# Thermosensitivity of the voltage-dependent activation of calcium homeostasis modulator 1 (calhm1) ion channel

**DOI:** 10.1101/2020.10.20.346940

**Authors:** Young Keul Jeon, Si Won Choi, Jae Won Kwon, Joohan Woo, Seong Woo Choi, Sang Jeong Kim, Sung Joon Kim

## Abstract

Calcium homeostasis modulator 1 (calhm1) proteins form an outwardly rectifying nonselective ion channel having exceedingly slow kinetics and low sensitivity to voltage that is shifted by lowering extracellular Ca^2+^ ([Ca^2+^]_e_). Here we found that physiological temperature dramatically facilitates the voltage-dependent activation of the calhm1 current (*I*_*calhm1*_); increased amplitude (Q_10_, 7-15) and fastened speed of activation. Also, the leftward shift of the half-activation voltage (*V*_*1/2*_) was similary observed in the normal and lower [Ca^2+^]_e_. Since calhm1 is highly expressed in the brain and taste cells, the thermosensitivity should be considered in their electrophysiology.

## Introduction

Calcium homeostasis modulator (calhm) family is comprised of several membrane proteins having four transmembrane (TM) domains with intracellular N- and C-termini. To distinguish the species, we denoted the human calhm as hCALHM and the mouse protein as calhm. The gene of human CALHM1 (*CALHM1*) was first discovered in a search for human genes linked to enhanced risk for late-onset Alzheimer’s disease [1]. Their multimeric assembly forms unselective ion channel with large pore structure, activated by membrane depolarization [1,2]. In mice, its knockout (*calhm1*^−/−^) impaired the long-term potentiation of hippocampal neurons [3], and a pathological role has been suggested in the ischemia-reperfusion injury of brain [4]. The expression of calhm1 has been also confirmed in taste buds, the nasal epithelium and the bladder, acting as a voltage-gated ATP-release channel [5–7]. hCALHM1 contains 346 amino acids comprising four TM domains and a long C-terminal structure containing four helix domains [2,8]. According to the recent cryo-electronmicroscopy studies, the hCALHM1 and hCALHM2 are composed in an octameric and undecameric assembly, respectively [9].

In the electrophysiological recordings of calhm1 at room temperature, even a slightly noticeable activation requires prolonged depolarization to above 10 mV with 82 mV of half-activation voltage (V_1/2_) [1,10,11]. Furthermore, the speed of activation was exceptionally slow, unable to reach a steady state within several seconds of the depolarized state [10,11]. Interestingly, the voltage-dependent activation of calhm1 or hCALHM1 is negatively modulated by extracellular Ca^2+^ concentration ([Ca^2+^]_e_). By decreasing [Ca^2+^]_e_, the voltage-dependence was shifted to the left, and the threshold voltage for activation was lowered [2,10].

The harsh conditions for activation, i.e. the low voltage-sensitivity and the slow activation kinetics under physiological ionic environment, cast doubt on to their physiological significance. Furthermore, in whole-cell patch clamp studies, the sustained clamp to high depolarization (e.g. > 40 mV) usually leads to unstable recordings in the mammalian cells, impeding precise electrophysiological studies.

Thermosensitivity is a key property of various ion channels, such as the vanilloid-type transient receptor potential channel (TRPV), Ca^2+^-activated Cl^−^ channels, voltage-dependent proton channels, and the TWIK-related K^+^ channel (TREK) [12–17]. Here, we investigated the effects of temperature on the voltage-dependence and kinetics of the calhm1 current (*I*_*calhm1*_) in human embryonic kidney cells (HEK293) cells using patch-clamp techniques. The present study revealed the remarkable facilitation of *I*_*calhm1*_ in the speed of activation as well as the voltage-dependence, at physiological temperatures.

## Materials and methods

### Cell culture and preparation

HEK293 cells were purchased from ATCC (Manassas, VA) and incubated in Dulbecco’s modified Eagle’s medium (DMEM; Gibco, Grand Island, NY) supplemented with 10% fetal bovine serum (FBS; Gibco) and 1% penicillin-streptomycin (Gibco). Cells were incubated at 37°C in 20% O_2_-5% CO_2_. Every day, the media was replaced with fresh media, and every 72 hours the cell were subcultured. The cells were centrifuged at 160 g for 2 min, and resuspended in fresh media. HEK293 cells at passages 5-15 were used for patch clamp experiments.

### Heterologous expression of calhms

The mouse or human complementary DNAs of *calhm1* (MR221396), *calhm2* (MR204675), *hCALHM1* (RC206902) and *hCLAHM3* (RC25514) were purchased from ORIGENE (Rockville, MD). All constructs were transfected into HEK293 cells using a TurboFect transfection reagent (ThermoFisher Scientific, Waltham, MA). The day before transfection, 1 × 10^5^ cells were seeded in a 12-well culture dish. The following day, 0.5-2.5 μg/well of pCMV6 vector containing the target cDNA was transfected into the cells. Twenty-four hours after transfection, the cells were passaged at a higher dilution (~50 cells/35-mm culture dish) into fresh medium.

### Electrophysiology

The cells were transferred to a bath mounted on the stage of an inverted microscope (TE20000-S; Nikon, Tokyo, Japan). The bath (0.15 ml) was continuously perfused at 5 ml·min^−1^. Borosilicated glass pipettes with a free-tip resistance of ~2.5 MΩ were connected to the CV 203Bu head stage of a patch-clamp amplifier (Axopatch 200B; Axon Instruments, San Jose, CA). The series resistance, estimated by dividing the time constants of capacitive current, was kept below 10 MΩ in the whole-cell configuration. To correct the cell size, the current amplitudes were divided by cell capacitance and expressed as pA·pF^−1^. The pipettes were pCLAMP software version 10.6.2 and Digidata-1440A (Axon Instruments) were used to acquire data and apply command pulses. The recorded currents were sampled at 10 kHz and were lowpass Bessel-filtered at 5 kHz.

### Temperature control

The temperatures of the bath and cells were controlled using the in-line solution heating system (Warner Instruments, Hamden, CT), and monitored by a thermistor in the bath near the cells (< 300 μm). The in-line solution heater was placed in front of bath (≅ 3 cm), and the heated perfusate was washed over the sample.

### Solutions and chemicals

Normal Tyrode’s bath solution (NT) comprised of (in mM) 140 NaCl, 5.4 KCl, 2 CaCl_2_, 1 MgCl_2_, 10 glucose, 20 mannitol, and 10 HEPES [4-(2-hydroxyethyl)-1-piperazine ethanesulfonic acid], with a pH of 7.4 (titrated with NaOH) was used for the whole-cell patch clamp experiments. The pipette solution for whole-cell patch clamp contained (in mM) 140 CsCl, 1 MgCl_2_, 10 EGTA [ethylene glycol-bis(β-aminoethyl ether)-*N,N,N’,N’*-tetra acetic acid], and 10 HEPES with a pH of 7.2 (titrated with CsOH). The chemicals and drugs used in this study were purchased from Sigma-Aldrich (St. Louis, MO).

### Data analysis

For the analysis of the activation speed, the normalized current-time data points of the different groups of were fitted to a single exponential function. The conductance of each recording was normalized to the maximum conductance (*G*_*max*_) and fitted by Boltzmann equation:

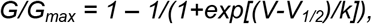

where *V*_*1/2*_ denotes the half maximum voltage (mV) and *k* is the slope factor (mV).

The time constant of activation, *τ*, was obtained by fitting the current trace with to a single exponential function:

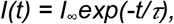

where *I*_∞_ is the estimated steady stat current amplitude and *t* is the time after the voltage step.

To assess the relative change in the current amplitude for a 10 °C change in temperature, the Q_10_ was calculated:

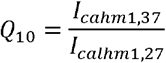

where *I*_*calhm1,37*_ is the current amplitude of calhm1 at 37°C and *I*_*calhm1,27*_ is that at 27°C.

### Data presentation and statistics

Data are presented as the mean ± standard deviation (S.D.). Curve fitting using the least squares method was performed in the Origin (Microcal Software) or handmade Python software. Student’s unpaired t-test was used where appropriate. Differences were considered as significant when *P* < 0.05.

## Results

Using the whole-cell patch clamp with the CsCl pipette solution, step-like depolarization from −20 to 40 mV (1 s), followed by repolarization to −40 mV (1 s) was applied at every 20 s. At 27°C, very small sizes of outward current (*I*_*calhm1*_) were induced by the depolarization. By raising the temperature of the bath perfusate to 37°C and 42°C, the amplitude of *I*_*calhm1*_ was markedly increased, showing a positive correlation between the bath temperature and the amplitude of *I*_*calhm1*_ (Fig. 1A, B). The *Q*_*10*_ value of *I*_*calhm1*_ at 40 mV was 13.5 ± 5.34 (n=25). It was also notable that the reversibility of the temperature effect was incomplete; when normalized to the maximum amplitude at 42°C, *I*_*calhm1*_ at the returned initial temperature (27°C) was 13.7 ± 2.15% (n=10), while *I*_*calhm1*_ at the same initial temperature (27°C) was 4.3 ± 0.57% of the maximum amplitude (Fig. 1C).

**Figure 1.**
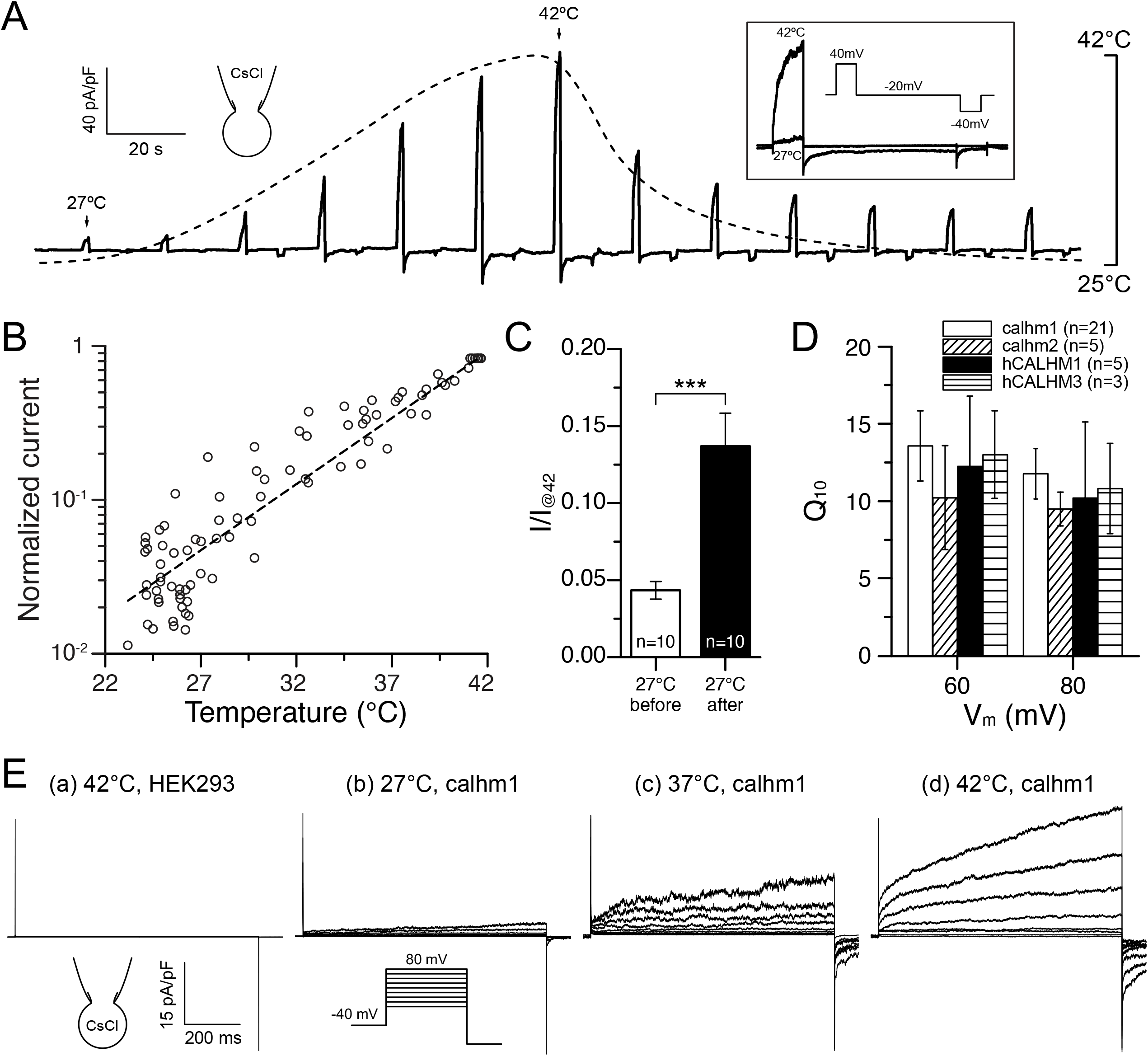
Thermosensitivity of calhm1 and CALHMs. (A) a representative chart trace of *I*_*calhm1*_ (black line) activated by repetitive step pulses from −20 to 40 mV (1 s). The bath temperature was concommitatnly recorded, and changed from 27 to 42°C (dotted line). *Inset* figure shows representative *I*_*calhm1*_ measured at 27 and 42°C, and the pulse protocol. (B) semi-logarithmic plot of the normalized amplitude of *I*_*calhm1*_ and bath temperature. From the data obtained from 12 cells as demonstrated in (A), the amplitude of *I*_*calhm1*_ was normalized to the maximum current at 42°C. The least-squares fitting of the data reveals a positive correlation between the bath temperature and *I*_*calhm1*_. (C) Normalized amplitudes of the *I*_*calhm1*_ at 27°C against 42°C (I/I_@42_), and their averaged summary comparing before and after the temperature change (n=10, \*\*\**P*<0.001). (D) Summary of the *Q*_*10*_ values at 60 and 80 mV in HEK293 cells overexpressing calhm1, calhm2, hCALHM1, and hCALHM3. (E) Representative traces of *I*_*calhm1*_ activated by incremental levels of step pulses (ranging from 10 to 80 mV, 1 s). The current amplitudes were increased at higher bath temperature.

Incremental levels of step pulses (ranging from 10 to 80 mV, 1 s) were applied from −40 mV of holding voltage (10 s interval between pulses), and the representative *I*_*calhm1*_ revealed slowly activating outward currents. The amplitudes of *I*_*calhm1*_ at 60 and 80 mV were similarly increased by raising the bath temperature from 27°C to 37°C or to 42°C (Fig. 1D, E). The temperature-dependent increase of the outward current was also observed in the HEK293 cells overexpressed with mouse calhm2, hCALHM1 and hCALHM3 (Fig. 1D). In the empty HEK293 cells, no significant level of the slowly activating outward current was observed even at 42°C (Fig. 1E), indicating that the temperature effect was not a nonspecific leak or artifact.

Then we analyzed the effect of temperature on the speed of *I*_*calhm1*_ development. To this end, a prolonged step-like depolarization (from −40 to 40 mV, 10 s) was applied at 27°C, 37°C and 42°C. The averaged traces of *I*_*calhm1*_ at the different temperaures demonstrate the acceleration of *I*_*calhm1*_ development (Fig. 2A). Notably, the temperature effect appeared to require a certain duration of depolarization. The ratio of I_calhm1_ amplitudes at 37°C and 42° (I/I_42°C_) were varied depending on the length of depolarization, and increased steeply between 0.5 and 1 s (Fig. 2B). Although an authentic steady-state was not obtained, the development of *I*_*calhm1*_ was fitted to a single exponential function (Fig. 2A, red dashed lines). The time constants (*τ*), which reflect the activation energy for channel opening, were significantly decreased when the temperature was increased (Fig. 2C). The *τ* at 40 mV were 466.6 ± 21.78 (n=27), 14.0 ± 4.01 (n=13), and 3.4 ± 0.49 s (n=16) at 27°C, 37°C and 42°C, respectively. When plotting the logarithm of the reciprocals of the time constant (*ln (1/τ)*) of *I*_*calhm1*_ at 40 mV against the bath temperature, a positive correlation was revealed (Fig. 2D). At each temperature tested, linear correlations were also observed between the time constants and the clamp voltages. This relationship was found to shift to the left by raising the temperature (Fig. 2D). These relationships implied that the activation energy for *I*_*calhm1*_ was a function of the temperature and membrane potential, respectively.

**Figure 2.**
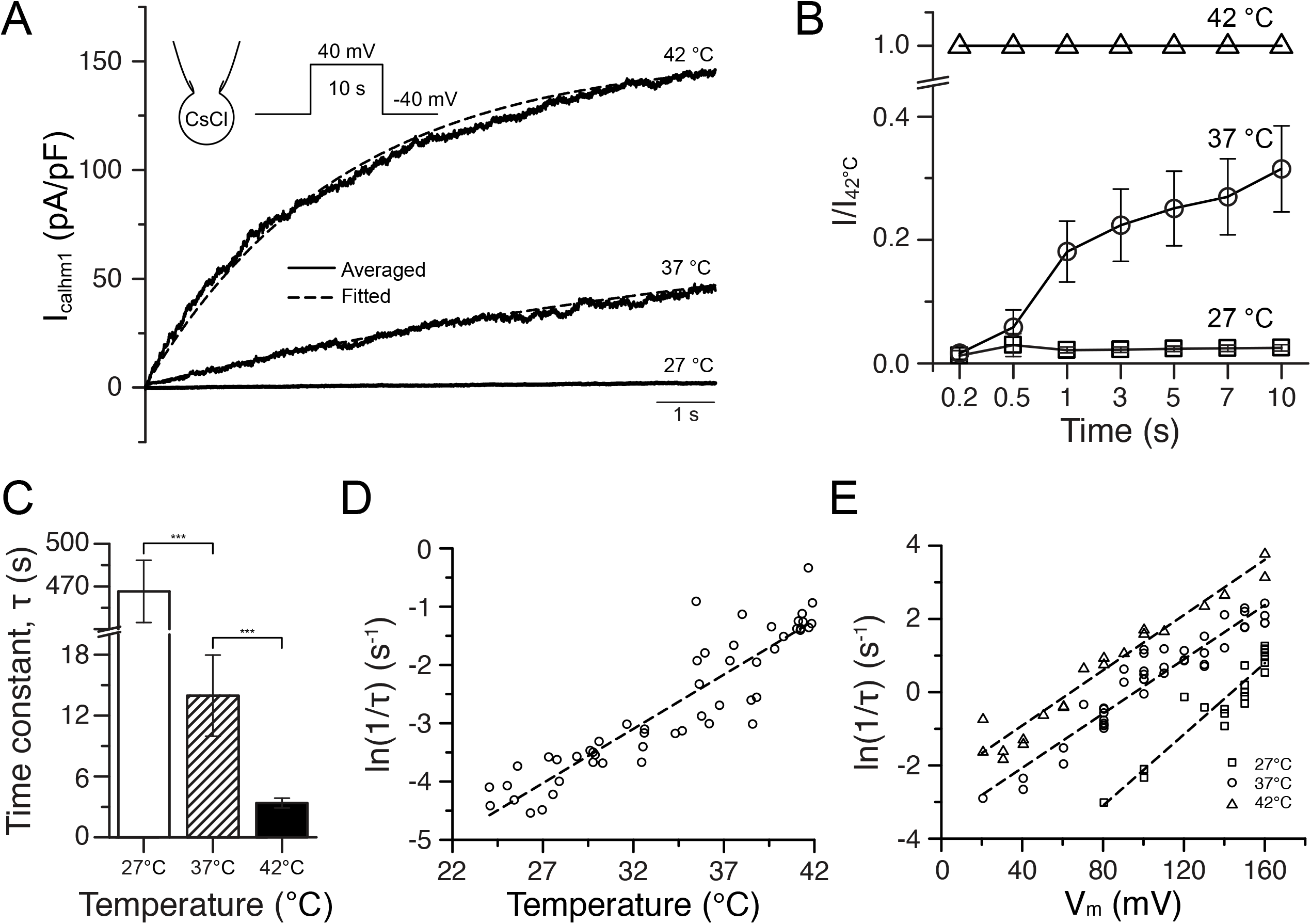
Effects of temperature on the speed of *I*_*calhm1*_ activation. (A) averaging traces of *I*_*calhm1*_ evoked by 10s of depolarization (40 mV) at three different temperatures (black line). Traces were fitted to single exponential function (red line). (B) Relative amplitudes of *I*_*calhm1*_ under 27°C and 37°C at different times of sustained depolarization normalized to the corresponding *I*_*calhm1*_ at 42°C (*I*/*I*_*@42*°C_). Steep increase was observed between 0.5 and 1 s. (C) Summary of the time constants at different temperatures at 27°C (n=22), 37°C (n=16) and 42°C (n=18). ***, *P*<0.001. (D, E) Arrhenius plots of the speed of *I*_*calhm1*_ activation at different temperature and membrane potentials. The logarithm of the reciprocals of the time constant (*ln (1/τ)*) of *I*_*calhm1*_ were affected by the bath temperature (D) as well as the membrane potentials (E).

Based on the relation between the pulse duration and the temperature effect (Fig. 2B), depolarizing step pulses with 1 s of duration were applied to analyze the voltage-dependence of *I*_*calhm1*_ and the effects of temperature (Fig. 3 and 4). It is known that the voltage-dependence of calhm1 is negatively affected by extracellular Ca^2+^([Ca^2+^]_e_); a lower level of [Ca^2+^]_e_ enables activation at a less depolarized state [8,10,11]. We measured the *I*_*calhm1*_ at 2, 1, and 0.5 mM of [Ca^2+^]_e_ with the multi-step pulse protocol. The facilitation by increased temperatures was still observed at a lower [Ca^2+^]_e_ (Fig. 3A). The current-to-voltage relation (I/V curve) became steeper and the threshold voltage of activation became lower with decreasing [Ca^2+^]_e_ from 2 to 1 and 0.5 mM (Fig. 3B-D).

**Figure 3.**
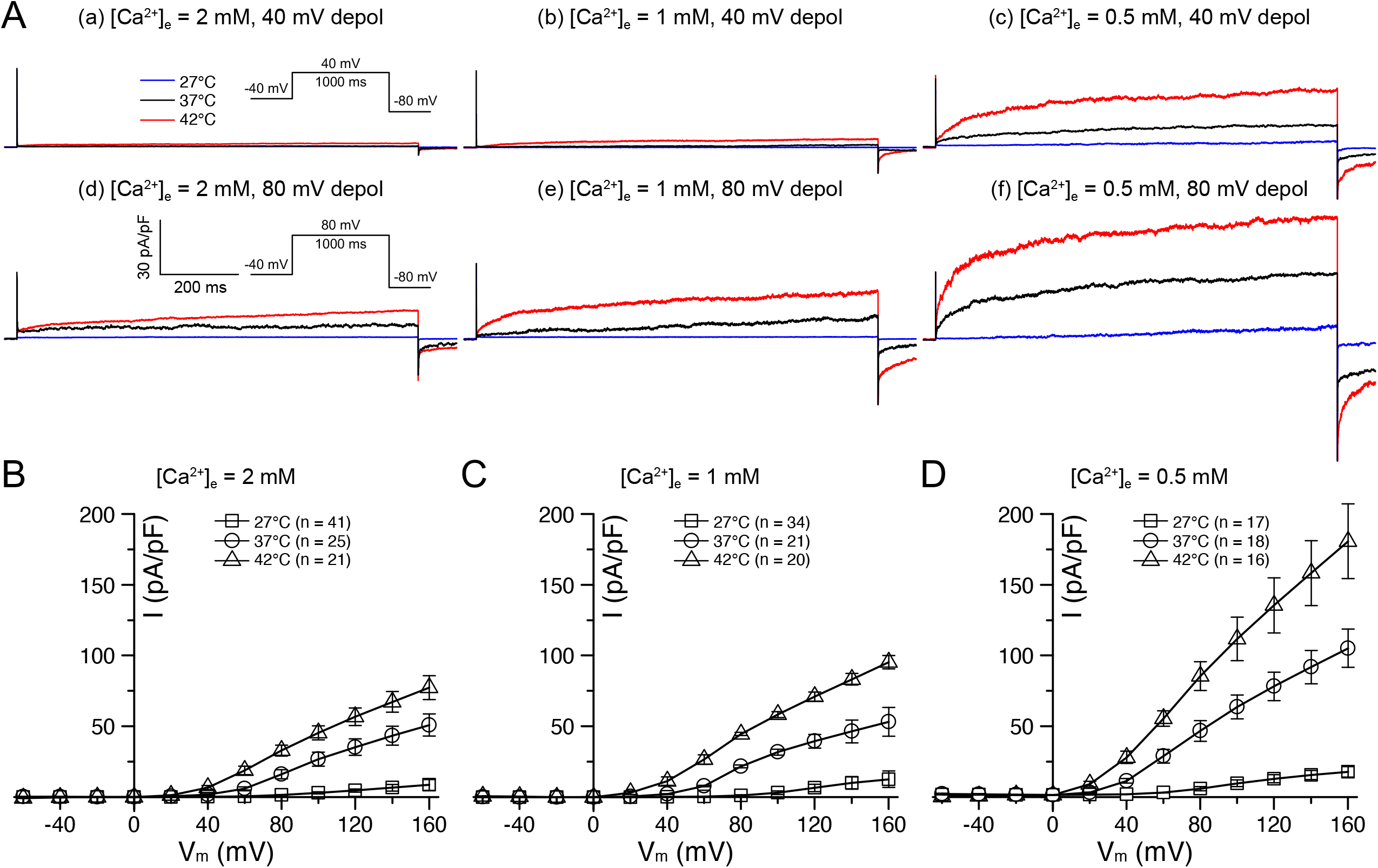
Voltage-dependence of *I*_*calhm1*_ and the effects of temperature and [Ca^2+^]_e_. (A) representative trace of *I*_*calhm1*_ at different [Ca^2+^]_e_ (2 mM, 1 mM, and 0.5 mM) and temperature (27°C (blue), 37°C (black), and 42°C (red)) conditions. *I*_*calhm1*_ were obtained by depolarization to 40 mV (upper panels) and 80 mV (lower panels). (B-D) current-to-voltage relations (I/V curves) of *I*_*calhm1*_ obtained by incremental levels of step pulse (ranging from −60 to 160 mV, 1 s) under different [Ca^2+^]_e_ and temperature (square; 27°C, circle; 37°C, and triangle; 42°C). [Ca^2+^]_e_ was 2 mM (B), 1 mM (C), and 0.5 mM (D).

**Figure 4.**
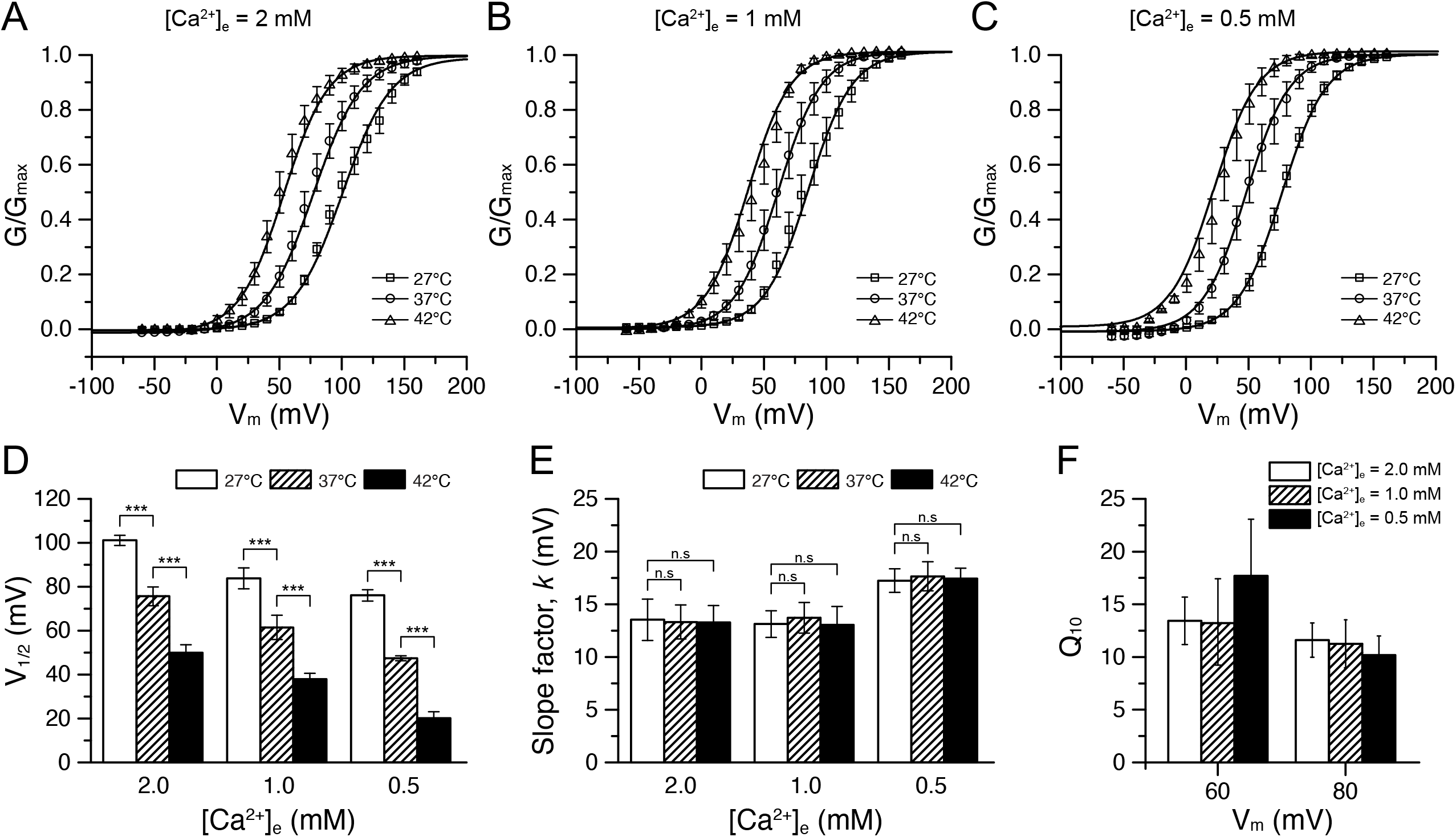
Analyses of the temperature- and [Ca^2+^]_e_ effects on the voltage dependence of *I*_*calhm1*_. (A-C) whole-cell conductance to voltage relations (G/V curves) of *I*_*calhm1*_ relationship under 2 mM (A), 1 mM (B), and 0.5 mM (C) of [Ca^2+^]_e_ and at different temperature. (D, E) summary of the half maximum activation voltages (*V*_*1/2*_) and slope factors (*k*) obtained by fitting the G/V curves to Boltzman equation. Decreases, i.e. leftward shift, of *V*_*1/2*_ by raised temperature was commonly observed at different [Ca^2+^]_e_ (D). In contrast, *k* was affected neither by [Ca^2+^]_e_ nor by temperature (E). (F) Effects of [Ca^2+^]_e_ on Q_10_ values. At the same voltage, the ratio of *I*_*calhm1,@37*°C_/ *I*_*calhm1,@27*°C_ (*Q*_*10*_) were not affected by the bath temperature.

To further analyze the effects of temperature on the voltage-dependence of calhm1, we determined the half-activation voltage (*V*_*1/2*_) and slope factor (*k*) obtained by fitting the conductance-to-voltage (G/V) curves of the channel with Boltzmann function (Fig. 4A-C). At 27°C in the NT solution, significant outward currents were only measured at highly depolarized voltages, with an estimated *V*_*1/2*_ of 102.2 ± 2.30 mV (n=41). At 37°C and 42°C, *V*_*1/2*_ values were 76.4 ± 4.32 (n=25) and 50.5 ± 3.72 (n=21), respectively (Fig. 4D). By lowering the [Ca^2+^]_e_, the *V*_*1/2*_ was also lowered. Furthermore, an increase in the temperature significantly shifted the *V*_*1/2*_ to the left, under 1 and 0.5 mM [Ca^2+^]_e_ (Fig. 4D). In contrast to the shift in *V*_*1/2*_, the slope factors were not affected by changing the temperature (Fig. 4E). The *Q*_*10*_ values of I_calhm1_ obtained at 1 and 0.5 mM [Ca^2+^]_e_ were not significantly different from those at 2 mM [Ca^2+^]_e_ (Fig. 4F).

## Discussion

The calhm (CALHM) is a relatively newly investigated membrane protein family forming unselective ion channels. Owing to the requirement of strong and sustained depolarization for their activation, their electrophysiological properties are poorly understood yet. Here, we firstly demonstrate the remarkable thermosensitivity of the mouse calhm1,-2 and the corresponding members of human CALHM ion channels. the amplitude of *I*_*calhm1*_ was also remarkably increased at the physiological temperature. Since the aqueous diffusion of ions and the reciprocal of the viscosity of water increase 1.4-fold after a 10 degree increase in the temperature, channels with a Q_10_ value higher than 5 are regarded as thermosensitive [18,19]. Since the averaged *Q*_*10*_ of calhm1, calhm2, hCALHM1 and hCALHM3 ranged from 10 to 13 (Fig. 1D), the calhm family could be safely classified as a thermosensitive channel. On depolarization, the speed of activation rose steeply by temperature, which could be fit to single exponential functions with shortened τ, indicating the thermosensitivity of the channel gating procedure (Fig. 2).

The voltage-dependence of calhm1 was also sensitized by temperature. As previously reported [10,11,20], *V*_*1/2*_ was left-shifted at a lower [Ca^2+^]_e_, and was further shifted by higher temperatures (Fig. 3). Although the precise mechanism for the thermosensitivity is yet unknown, the additive effects of lower [Ca^2+^]_e_ and the higher temperature suggest independent mechanisms for the modulation of voltage-sensitivity.

Recently, cryo-electron microscopic investigations have revealed the molecular structure of the calhm channels [9,20,21]. In contrast to the undecameric assembly of hCALHM2, the structure of hCALHM1 shows an octameric organization with a smaller pore diameter [20–22], which suggest that the functional properties of activation and conductance may not be conserved within the family. However, the results of the present study show that the thermosensitivity is conserved between the tested paralogs including hCALHM3.

### Physiological implication

Despite the shift of *V*_*1/2*_, the raised temperature alone could not evoke inward current at the negative holding potential (e.g. −40 mV), which is different from the classical thermo-activating channels such as TRPV1. Nevertheless, the slow kinetics of deactivation on the repolarization and the long-lasting effect of high temperature (Fig. 1C, E) suggest that a cumulative activation of calhm might affect the intrinsic escitability of the cells expressing the channels. The plausible effects of temperature could be expected in the excitable cells generating action potentials (AP). In the hippocampal and cortical neurons, the genetic knock-out of calhm1 affected their excitability [3,10]. Furthermore, the [Ca^2+^]_e_ of diffusion-limited intercellular spaces or synaptic cleft could be significantly lowered depending on the Ca^2+^ influx-associated AP firing [23]. In addition to the spatial fluctuation of [Ca^2+^]_e_, the present study warrants to consider the temperature effect on the roles of calhm1 *in vivo*.

Calhm1 channels are permeable not only to cations but also to various sizes of anions, including ATP^4−^ (ATP). The ATP-permeability is thought to be responsible for the purinergic signaling between the taste receptor cells and sensory nerve endings, a process known as non-vesicular type synaptic transmission [5]. The calhm channels (calhm1 and 3) in the taste cells could be activated during the burst of APs [5,6,22]. Previous studies have reported on the temperature-dependent facilitation of some taste transduction efficiencies [24,25]. In addition to the warmth-activated TRPM5 cation channel in the gustatory cells [5,25,26], the thermosensitivity of calhm may partly explain the effects of temperature on the taste sensation. Another potential contribution of thermosensitive calhm is in the ciliary beating of the airway epithelium, where the release of ATP via calhm1 may control the ciliary beating frequency (CBF) [7]. CBF in airway is known to be markedly increased by rises in temperature from room temperature to physiological temperatures [27,28].

Calhm2 has been suggested to play a critical role as an ATP-releasing pathway has been also suggested in the mouse astrocytes; calhm2 mediates ATP release and the knockout of calhm2 leads to the neural dysfunction and depression-like phenotypes in mice [29]. As calhm2 was confirmed to exhibit thermosensitivity in our present study, it is likely that the release of ATP from astrocytes may be also boosted at physiological temperatures.

In summary, the effective temperature-dependent facilitation of the calhm1 activation was demonstrated. Our findings provide a basis for further studies with the aim of elucidating their electrophysiological properties. Also, the thermosensitivity demonstrated in the present study provides insights that could broaden the physiological understanding of the roles of calhm family channels *in vivo*.

## Acknowledgements

This work was supported by National Research Foundation of Korea (NRF-2018R1A5A2025964) and by the 2020 Research Fund from Seoul National University Hospital to Sung Joon K.

## Author Contributions

Young Keul J., Seong Woo C., Sang Jeong K., and Sung Joon K. designed the research; Young Keul J., Jae Won K., Si Won C., and Joo Han W. performed experiments; Young Keul J., and Jae Won K. analyzed data; Si Won C. and Seong Woo C. interpreted data; Young Keul J. prepared figures; Young Keul J. drafted manuscripts; Sung Joon K. and Sang Jeong K. edited revised manuscripts.

## References

[1] U. Dreses-Werringloer, J.C. Lambert, V. Vingtdeux, H. Zhao, H. Vais, A. Siebert, A. Jain, J. Koppel, A. Rovelet-Lecrux, D. Hannequin, F. Pasquier, D. Galimberti, E. Scarpini, D. Mann, C. Lendon, D. Campion, P. Amouyel, P. Davies, J.K. Foskett, F. Campagne, P. Marambaud, A polymorphism in CALHM1 influences Ca2+ homeostasis, Abeta levels, and Alzheimer’s disease risk, Cell 133 (2008) 1149–1161. 10.1016/j.cell.2008.05.048.

[2] Z. Ma, J.E. Tanis, A. Taruno, J.K. Foskett, Calcium homeostasis modulator (CALHM) ion channels, Pflugers Arch 468 (2016) 395–403. 10.1007/s00424-015-1757-6.

[3] V. Vingtdeux, E.H. Chang, S.A. Frattini, H. Zhao, P. Chandakkar, L. Adrien, J.J. Strohl, E.L. Gibson, M. Ohmoto, I. Matsumoto, P.T. Huerta, P. Marambaud, CALHM1 deficiency impairs cerebral neuron activity and memory flexibility in mice, Sci Rep 6 (2016) 24250. 10.1038/srep24250.

[4] J. Garrosa, I. Paredes, P. Marambaud, M.G. Lopez, M.F. Cano-Abad, Molecular and pharmacological modulation of CALHM1 promote neuroprotection against oxygen and glucose deprivation in a model of hippocampal slices, Cells 9 (2020). 10.3390/cells9030664.

[5] A. Taruno, V. Vingtdeux, M. Ohmoto, Z. Ma, G. Dvoryanchikov, A. Li, L. Adrien, H. Zhao, S. Leung, M. Abernethy, J. Koppel, P. Davies, M.M. Civan, N. Chaudhari, I. Matsumoto, G. Hellekant, M.G. Tordoff, P. Marambaud, J.K. Foskett, CALHM1 ion channel mediates purinergic neurotransmission of sweet, bitter and umami tastes, Nature 495 (2013) 223–226. 10.1038/nature11906.

[6] Z. Ma, W.T. Saung, J.K. Foskett, Action potentials and ion conductances in wild-type and CALHM1-knockout type II taste cells. J Neurophysiol 117 (2017) 1865–1876. 10.1152/jn.00835.2016.

[7] A.D. Workman, R.M. Carey, B. Chen, C.J. Saunders, P. Marambaud, C.H. Mitchell, M.G. Tordoff, R.J. Lee, N.A. Cohen, CALHM1-Mediated ATP Release and Ciliary Beat Frequency Modulation in Nasal Epithelial Cells. Sci Rep 7 (2017) 6687. 10.1038/s41598-017-07221-9.

[8] A.P. Siebert, Z. Ma, J.D. Grevet, A. Demuro, I. Parker, J.K. Foskett, Structural and functional similarities of calcium homeostasis modulator 1 (CALHM1) ion channel with connexins, pannexins, and innexins. J Biol Chem 288 (2013) 6140–6153. 10.1074/jbc.M112.409789.

[9] W. Choi, N. Clemente, W. Sun, J. Du, W. Lu, The structures and gating mechanism of human calcium homeostasis modulator 2, Nature 576 (2019) 163–167. 10.1038/s41586-019-1781-3.

[10] Z. Ma, A.P. Siebert, K.H. Cheung, R.J. Lee, B. Johnson, A.S. Cohen, V. Vingtdeux, P. Maram baud, J.K. Foskett, Calcium homeostasis modulator 1 (CALHM1) is the pore-forming subunit of an ion channel that mediates extracellular Ca2+ regulation of neuronal excitability. Proc Natl Acad Sci U S A 109 (2012) E1963–1971. 10.1073/pnas.1204023109.

[11] J.E. Tanis, Z. Ma, J.K. Foskett, The NH2 terminus regulates voltage-dependent gating of CAL HM ion channels. Am J Physiol Cell Physiol 313 (2017) C173–C186. 10.1152/ajpcell.00318.2016.

[12] C. Arrigoni, D.L. Minor, Jr., Global versus local mechanisms of temperature sensing in ion channels, Pflügers Arch 470 (2018) 733–744. 10.1007/s00424-017-2102-z.

[13] M.R. Cohen, V.Y. Moiseenkova-Bell, Structure of thermally activated TRP channels. Curr Top Membr 74 (2014) 181–211. 10.1016/B978-0-12-800181-3.00007-5.

[14] I. Diaz-Franulic, H. Poblete, G. Mino-Galaz, C. Gonzalez, R. Latorre, Allosterism and Structure in Thermally Activated Transient Receptor Potential Channels. Annu Rev Biophys 45 (2016) 371–398. 10.1146/annurev-biophys-062215-011034.

[15] D. Kang, C. Choe, D. Kim, Thermosensitivity of the two-pore domain K+ channels TREK-2 an d TRAAK. J Physiol 564 (2005) 103–116. 10.1113/jphysiol.2004.081059.

[16] H. Lin, I. Jun, J.H. Woo, M.G. Lee, S.J. Kim, J.H. Nam, Temperature-dependent increase in the calcium sensitivity and acceleration of activation of ANO6 chloride channel variants. Sci Rep 9 (2019) 6706. 10.1038/s41598-019-43162-1.

[17] F. Maingret, I. Lauritzen, A.J. Patel, C. Heurteaux, R. Reyes, F. Lesage, M. Lazdunski, E. Honore, TREK-1 is a heat-activated background K+ channel, EMBO J 19 (2000) 2483–2491. 10.1093/emboj/19.11.2483.

[18] B. Frankenhaeuser, L.E. Moore, The effect of temperature on the sodium and potassium permeability changes in myelinated nerve fibres of Xenopus Laevis. J Physiol 169 (1963) 431–437. 10.1113/jphysiol.1963.sp007269.

[19] K.G. Beam, P.L. Donaldson, A quantitative study of potassium channel kinetics in rat skeletal muscle from 1 to 37°C, J Gen Physiol 81 (1983) 485–512. 10.1085/jgp.81.4.485.

[20] J.L. Syrjanen, K. Michalski, T.H. Chou, T. Grant, S. Rao, N. Simorowski, S.J. Tucker, N. Grigorieff, H. Furukawa, Structure and assembly of calcium homeostasis modulator proteins. Nat Struct Mol Biol 27 (2020) 150–159. 10.1038/s41594-019-0369-9.

[21] K. Drozdzyk, M. Sawicka, M.I. Bahamonde-Santos, Z. Jonas, D. Deneka, C. Albrecht, R. Dutzler, Cryo-EM structures and functional properties of CALHM channels of the human placenta, Elife 9 (2020). 10.7554/eLife.55853.

[22] K. Nomura, M. Nakanishi, F. Ishidate, K. Iwata, A. Taruno, All-Electrical Ca2+-Independent Signal Transduction Mediates Attractive Sodium Taste in Taste Buds, Neuron 106 (2020) 816–829 e816. 10.1016/j.neuron.2020.03.006.

[23] D.A. Rusakov, A. Fine, Extracellular Ca2+ depletion contributes to fast activity-dependent modulation of synaptic transmission in the brain, Neuron 37 (2003) 287–297. 10.1016/s0896-6273(03)00025-4.

[24] B.G. Green, C. Alvarado, K. Andrew, D. Nachtigal, The effect of temperature on umami taste. Chem Senses 41 (2016) 537–545. 10.1093/chemse/bjw058.

[25] C.H. Lemon, Modulation of taste processing by temperature. Am J Physiol Regul Integr Comp Physiol 313 (2017) R305–R321. 10.1152/ajpregu.00089.2017.

[26] K. Talavera, K. Yasumatsu, T. Voets, G. Droogmans, N. Shigemura, Y. Ninomiya, R.F. Margolskee, B. Nilius, Heat activation of TRPM5 underlies thermal sensitivity of sweet taste, Nature 438 (2005) 1022–1025. 10.1038/nature04248.

[27] C.F. Clary-Meinesz, J. Cosson, P. Huitorel, B. Blaive, Temperature effect on the ciliary beat frequency of human nasal and tracheal ciliated cells. Biol Cell 76 (1992) 335–338. 10.1016/0248-4900(92)90436-5.

[28] A. Green, L.A. Smallman, A.C. Logan, A.B. Drake-Lee, The effect of temperature on nasal ciliary beat frequency. Clin Otolaryngol Allied Sci 20 (1995) 178–180. 10.1111/j.1365-2273.1995.tb00040.x.

[29] M. Jun, Q. Xiaolong, Y. Chaojuan, P. Ruiyuan, W. Shukun, W. Junbing, H. Li, C. Hong, C. Jinbo, W. Rong, L. Yajin, M. Lanqun, W. Fengchao, W. Zhiying, A. Jianxiong, W. Yun, Z. Xia, Z. Chen, Y. Zengqiang, Calhm2 governs astrocytic ATP releasing in the development of depression-like behaviors. Mol Psychiatry 23 (2018) 1091. 10.1038/mp.2017.254.

